# The attenuation of activity-based anorexia by obese adipose tissue transplant is AgRP neuron-dependent

**DOI:** 10.1101/2024.04.23.590824

**Authors:** Dongmin J. Yoon, Jie Zhang, Rizaldy C. Zapata, Martina Ulivieri, Avraham M. Libster, Matthew S. McMurray, Olivia Osborn, Stephanie C. Dulawa

## Abstract

Anorexia nervosa (AN) is an eating disorder observed primarily in girls and women, and is characterized by a low body mass index, hypophagia, and hyperactivity. The activity-based anorexia (ABA) paradigm models aspects of AN, and refers to the progressive weight loss, hypophagia, and hyperactivity developed by rodents exposed to time-restricted feeding and running wheel access. Recent studies identified white adipose tissue (WAT) as a primary location of the ‘metabolic memory’ of prior obesity, and implicated WAT-derived signals as drivers of recidivism to obesity following weight loss. Here, we tested whether an obese WAT transplant could attenuate ABA-induced weight loss in normal female mice. Recipient mice received a WAT transplant harvested from normal chow-fed, or HFD-fed obese mice; obese fat recipient (OFR) and control fat recipient (CFR) mice were then tested for ABA. During ABA, OFR mice survived longer than CFR mice, defined as maintaining 75% of their initial body weight. Next, we tested whether agouti-related peptide (AgRP) neurons, which regulate feeding behavior and metabolic sensing, mediate this effect of obese WAT transplant. CFR and OFR mice received either control or neonatal AgRP ablation, and were assessed for ABA. OFR intact mice maintained higher body weights longer than CFR intact mice, and this effect was abolished by neonatal AgRP ablation; further, ablation reduced survival in OFR, but not CFR mice. In summary, obese WAT transplant communicates with AgRP neurons to increase body weight maintenance during ABA. These findings encourage the examination of obese WAT-derived factors as potential treatments for AN.

## INTRODUCTION

Anorexia nervosa (**AN**) is a complex illness mostly affecting women and girls (1), and is characterized by hypophagia, dangerously low body weight, and compulsive exercise (2–5). AN has a lifetime prevalence of approximately 1%, and has one of the highest mortality rates of all psychiatric disorders (1). Although numerous classes of therapeutics have been examined in AN including antipsychotics, antidepressants, and anti-epileptics, no treatment has been consistently shown to have long-term efficacy for weight gain in this patient population (6–10). A recent large-scale genetic analysis reported that AN shows significant genetic correlations with metabolic traits, and should be reconceptualized as a metabo-psychiatric disorder (11, 12). Still, no agents specifically targeting metabolic processes have been explored as therapeutics for AN.

Activity-based anorexia (ABA) is a phenomenon observed ubiquitously in normal rodents and other mammals (13–15), and recapitulates several aspects of AN (16–21). In the ABA paradigm, rodents are subjected to time-scheduled feeding and have continuous access to running wheels. Under these conditions, animals develop rapid and progressive weight loss, hypophagia, and compulsive wheel running. Contrary to this, animals exposed to either time-scheduled feeding or constant running wheel access maintain body weight indefinitely (16, 22). Thus, in the ABA paradigm, animals with negative energy balance paradoxically choose to exercise more and reduce food intake, even during times of food availability. If allowed to continue unchecked, the ABA paradigm results in hypothermia, increased HPA axis activity, and ultimately stomach ulceration and death (16, 23, 24). Younger animals lose body weight more rapidly than older animals during ABA (25), paralleling clinical observations that AN is a disorder with peak onset and prevalence during adolescence (26, 27).

White adipose tissue (WAT) plays an active role in the regulation of energy balance as an endocrine organ (28–31). Weight loss through dieting is frequently followed by weight regain due to compensatory changes that encourage a higher body weight (30–33). Body weight recidivism following dieting was recently suggested to be driven by a “metabolic memory” of obesity, which predominantly resides in WAT (28, 34). Specifically, gene expression and metabolites within metabolic tissues were compared between lean, obese, or previously obese mice; obesity induced widespread transcriptional and metabolomic changes that were reversible by weight loss across most metabolic tissues, except for WAT (34). Thus, WAT-driven metabolic memory of obesity has been implicated in weight regain. Furthermore, crosstalk between WAT and brain regions including the hypothalamus is mediated by the sympathetic nervous system and circulating factors secreted by WAT (35, 36). Identifying the interactions between WAT and the brain that modulate body weight could lead to novel treatments for AN.

Agouti-related peptide (AgRP)-expressing neurons are localized in the arcuate nucleus (ARC) of the hypothalamus and regulate metabolism, food intake, and complex non-feeding behaviors that promote food pursuit (37–40). For example, AgRP neurons integrate internal signals of energy state with external signals relating to energy state, such as food availability and environmental cues of energy demand, to drive foraging (41–43). Recently, neonatal AgRP ablation was reported to increase ABA severity (44), while chemogenetic activation of AgRP neurons reduced ABA progression (44, 45). Furthermore, activated AgRP neurons shift metabolism towards energy conservation and lipid storage (37, 46–49). Here, we hypothesized that AgRP neurons would receive cues from obese WAT transplants that would alter the metabolic and/or behavioral state of recipient mice to increase body weight maintenance during ABA.

We determined whether transplanting perigonadal WAT from obese HFD-fed mice into normal weight recipient mice would attenuate ABA in the recipients. First, we compared the effects of control versus obese WAT transplant on survival, body weight, food intake, and wheel running in the ABA paradigm. Second, we tested whether neonatal ablation of AgRP-expressing neurons attenuates the ability of obese WAT transplant to reduce ABA-induced weight loss. We used young adult female mice, since AN predominantly affects young women and girls (1, 50).

## METHODS

### Mice

All mice were housed in a climate-controlled room with food and water available *ad libitum* unless otherwise stated. For Experiment 1, mice were maintained on a 12:12 light–dark cycle (lights on at 0700). Weeks before Experiment 2 was conducted, the mouse colony room was switched to a reversed 12:12 light–dark cycle (lights off at 0700). All procedures were approved by the Institutional Animal Care and Use Committee at University of California, San Diego.

All donor and recipient mice were young adult female wild type (WT) C57BL/6J mice purchased from Jackson (JAX) Laboratories. For Experiment 2, donor mice were WT C57BL/6J and recipient mice were AgRP-iCre^iDTR+/-^ knockins bred in house. Heterozygous mice expressing Cre recombinase under the control of the AgRP promotor: AgRP-iCre^+/-^ mice, strain AgRP^tm1(Cre)Low1/J^ (012899, JAX) were bred with mice which were homozygous for a floxed and inactivated simian diphtheria toxin receptor: iDTR mice, strain C57BL/6-Gt(ROSA)26Sor^tm1(HBEGF)Awai/J^ (007900, JAX). Thus, all experimental offspring carried one iDTR allele, plus either a control or an AgRP-iCre allele. Consequently, about half of the experimental offspring had AgRP neurons that were susceptible to ablation following diphtheria toxin administration.

### AgRP neuron ablation

In Experiment 2, all experimental offspring were injected subcutaneously with diphtheria toxin (50μg/kg) from between P5-P7, as previously described (44, 51). Diphtheria toxin was dissolved in phosphate-buffered saline (PBS; 13.3 µl/g body weight). Thus, diptheria toxin injections resulted in mice with ablated AgRP neurons (AgRP-iCre^iDTR+/-^) or intact AgRP neurons (WT^iDTR+/-^)(Figure 1).

**Figure 1.**
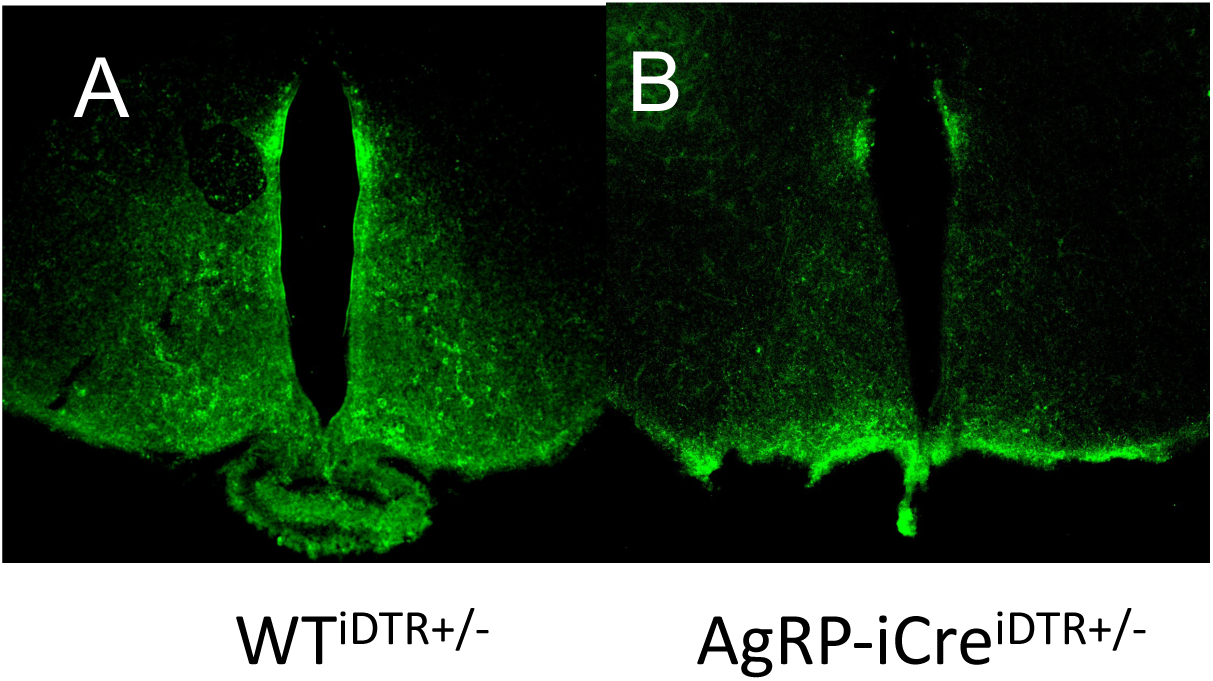
Ablation of AgRP neurons by neonatal diphtheria toxin injections. Both control and AgRP-iCre ^DTR+/-^ mice received injections as pups with diphtheria toxin (50ug/kg). Representative immunostaining of ARC neurons of (A) control, and (B) AgRP-iCre ^DTR+/-^ mice during early adulthood.

### Adipose tissue transplant

All experimental mice received WAT transplants from either normal weight or obese donors (Figure 2a). Beginning at 10 weeks of age, donor mice were fed either HFD (60% calories from fat, Research Diets) to develop obesity, or normal chow. Donors were sacrificed at 19 weeks of age, and intra-abdominal perigonadal WAT was cut into 200 mg slices for obese WAT, or 100 mg for normal weight WAT, and kept in saline-filled 50 ml tubes in a 37°C water bath until transplantation. Double the weight of obese WAT was transplanted compared to normal weight WAT to ensure that equal number of cells were transplanted from both groups, since adipocytes in the obese state are twice as large as normal adipocytes (52). At 10-12 weeks of age, obese fat recipients (OFR) received obese WAT transplants, while control fat recipient mice (CFR) received WAT from normal weight donors (Figure 2b). Recipient mice were anesthetized, and donor WAT slices were sutured next to the mesenteric fat below the liver to increase vascular support. After a 4-week recovery, recipient mice were tested in the ABA paradigm.

**Figure 2.**
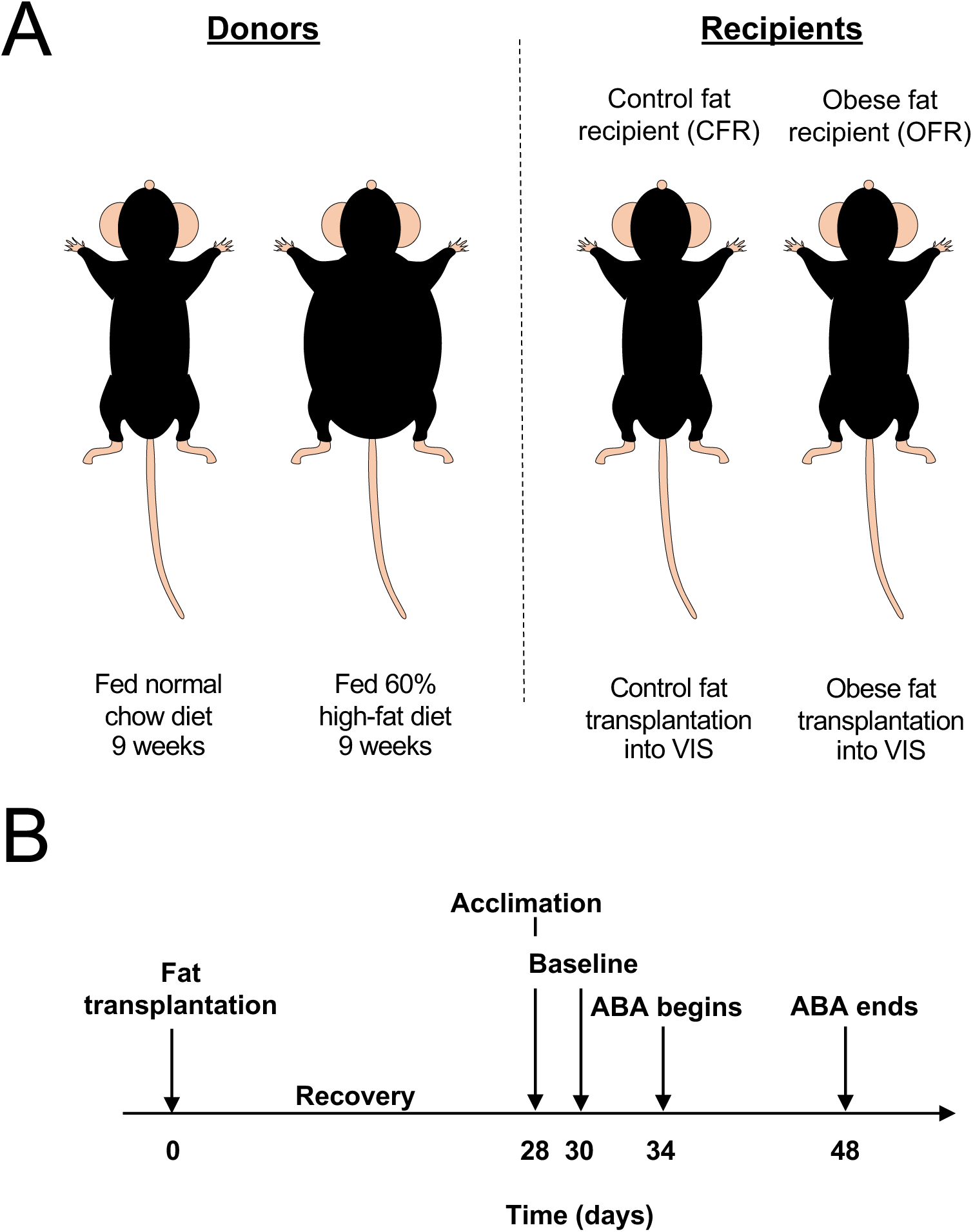
Experimental design. (A) Recipient mice received WAT from either high fat diet (HFD)-fed obese mice or normal chow-fed donor mice. Control or obese WAT was transplanted into the visceral area of normal weight recipient mice. (B) Acclimation to ABA housing conditions began 4 weeks after WAT transplantation. After 2 days of acclimation, 4 days of baseline testing occurred, followed by 2 weeks of testing during restriction conditions.

### Activity-Based Anorexia Paradigm

For both experiments, recipient mice 14-16 weeks of age were singly housed in home cages (19.56 × 34.70 × 14.41 cm). Each cage was equipped with a wireless low-profile running wheel (Med Associates, St Albans, VT, USA), which continuously transmitted running data every 30 s, 24 h a day to a computer with Wheel Manager software. Equipment failure resulted in loss of running wheel data on restriction day 9 of Experiment 2. Food (standard chow) was provided *ad libitum* in a glass jar (5 cm diameter × 4 cm height) resting on the cage floor, and tap water was available *ad libitum* in standard bottles.

Mice were acclimated for 2 days to single housing in the experimental cage. Then, mice entered the baseline phase (4 days), during which standard chow was constantly available, and daily body weight, food intake, and wheel running data were collected. Next, all mice entered the restriction phase, during which food was available for only for 3 h a day beginning at 0900 h. During restriction, mice “drop out” of the paradigm once they lose 25% of their initial body weight which was measured on day 4 of the baseline phase. Mice were sacrificed immediately following drop out, or when the study ended at day 14 (Figure 2b). Body weight, food intake, and wheel running were recorded daily during both baseline and restriction conditions. The experimenter was blinded to the groups.

For Experiment 1, CFR (n=21) and OFR (n=21) mice were assessed in the ABA paradigm (Figure 2). In Experiment 2, CFR and OFR mice additionally underwent either control or neonatal AgRP ablation; thus, four groups of animals were assessed for ABA: CFR intact (n=12), CFR ablated (n=14), OFR intact (n=12), and OFR ablated (n=13).

### Data Analysis

During the restriction phase of ABA, the number of days required for mice to meet the criterion for drop out (loss of 25% of baseline day 4 body weight) was analyzed using Cox proportional-hazards model (survival analysis). Analysis of the behavioral measures collected during ABA creates statistical challenges due to the presence of expected missing values in the restricted phase, as animals drop out. Thus, general linear mixed models (proc glimmix; SAS v9.2) were used to assess baseline and restriction outcomes, including factors for day, transplant group (control versus obese), and AgRP ablation group (intact versus ablated) for Experiment 2. Separate analyses were performed for body weight, food intake, and wheel running. Food anticipatory activity (FAA, wheel running during the 4-hour period before food delivery), and postprandial activity (PPA, wheel running during the period after food removal until lights off at 1900h) were analyzed in the same fashion. When significant main effects or interactions were observed, post-hoc tests were applied and adjusted for multiple comparisons using the Tukey-Kramer method. Alpha levels were set at p<0.05.

## RESULTS

### Obese WAT transplant reduces weight-loss in the ABA paradigm

In Experiment 1, no difference in body weight was found between CFR and OFR mice; however, a main effect of day was observed (F(_3,120_)= 4.72, p<0.01; Figure 3a), with a small increase in body weight emerging over days. Similarly, there was no difference in food intake or wheel running between CFR and OFR mice during baseline (Figure 3b,c). Additionally, main effects of day were found for food intake (F(_3,120_)= 2.85, p<0.05; Figure 3b) and wheel running (F(_3,114_)= 5.29, p<0.01; Figure 3c) reflecting small increases in these measures over days. Overall, no differences in body weight, food intake, or wheel running were found between CFR versus OFR mice during baseline.

**Figure 3.**
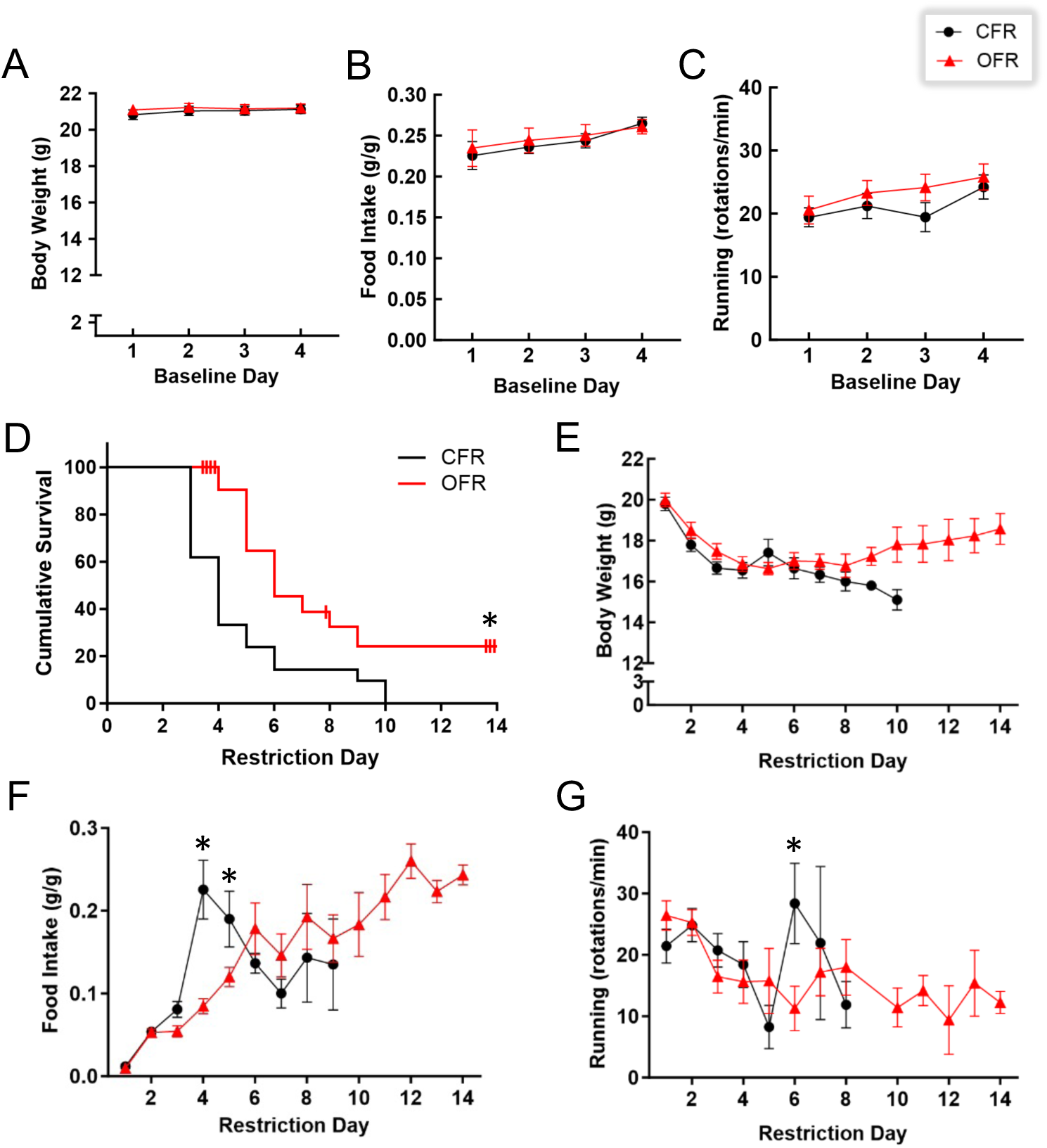
Experiment 1: Effects of obese WAT transplant on ABA. (A) Body weight in grams of CFR and OFR mice over the 4 days. (B) Food intake in grams divided by body weight of CFR and OFR mice over the 4 days. (C) Wheel running revolutions of CFR and OFR mice over the 4 days. (D) OFR mice survived significantly longer than CFR mice during restriction. (E) Body weight of CFR and OFR mice over the 14 days of restriction. (F) CFR mice ate significantly more than CFR mice on day 4 of restriction. (G) Wheel running of CFR and OFR mice over the 14 days of restriction. Data are adjusted mean values ± SEM, n = 21/group. An asterisk (*) indicates a significant effect of transplant group.

Interestingly, during the restriction period of Experiment 1, OFR mice survived longer in the paradigm than CFR mice (Χ^2^=7.51, p<0.01; Figure 3d). Hazard ratios indicated that CFR mice were 2.72 times more likely to drop out of the study than OFR mice (95% CI: 1.338-5.596). The longer survival of OFR compared to CFR mice reflects that OFR mice required a longer time to reach the 25% weight loss criterion; thus, survival provides one measure of body weight. In sum, obese WAT transplant increased survival under ABA conditions.

Although survival differed between the two transplant groups (Figure 3d), body weight values for animals still in the study did not differ between CFR and OFR mice during restriction (Figure 3e). For food intake, a two-way interaction of transplant group and day (F(_8, 137_)= 8.84, p<0.001) and post-hoc tests revealed that CFR mice showed a transient increase in consumption relative to OFR mice on day 4 (p<0.001) and day 5 (p<0.001) of restriction (Figure 3f). This sharp increase in food intake by CFR mice coincided with a large number of CFR mice dropping out of the paradigm on days 3 and 4 (Figure 3d). This transient increase in food intake might reflect the higher food intake values of remaining CFR mice.

For wheel running, there was a two-way interaction of transplant group and day (F(_7, 128_)= 2.29, p<0.05). Post-hoc tests revealed that CFR mice showed more running relative to OFR mice on day 6 of restriction (p<0.01)(Figure 3g). It should be noted that in survival studies, the number of mice in the study decreases over time, which can result in a large standard error of the mean as days progress, such as for the CFR group with only 3 mice remaining on day 7 (Figure 3g). To determine whether the transplant groups differed in wheel running during specific phases of ABA, we analyzed wheel running during the FAA and PPA periods. Although main effects of day were observed for both FAA (F(_11,127_)= 2.88, p<0.01; Supplemental Figure 1a) and PPA (F(_13,131_)= 3.81, p<0.001; Supplemental Figure 1b), no main effects or interactions were found for wheel running during the FAA or PPA period.

### AgRP neurons are required for obese WAT transplant to increase body weight during ABA

In Experiment 2, OFR mice showed a small but significant increase in body weight compared to CFR mice across days during baseline (F(_1, 47_)= 8.90, p<0.01; Figure 4a,b). Further, a main effect of day was observed, with body weight increasing over baseline days (F(_3, 141_)= 26.70, p<0.001; Figure 4a). No difference in food intake or wheel running was found between transplant groups (Figure 4b,c). Main effects of day were found for both food intake (F(_3,141_)= 22.87, p<0.001; Figure 4b) and wheel running (F(_3,129_)= 9.60, p<0.001; Figure 4c), reflecting small differences over days. Overall, CFR and OFR groups differed only in body weight during baseline, with OFR mice showing slightly higher body weights than CFR mice.

**Figure 4.**
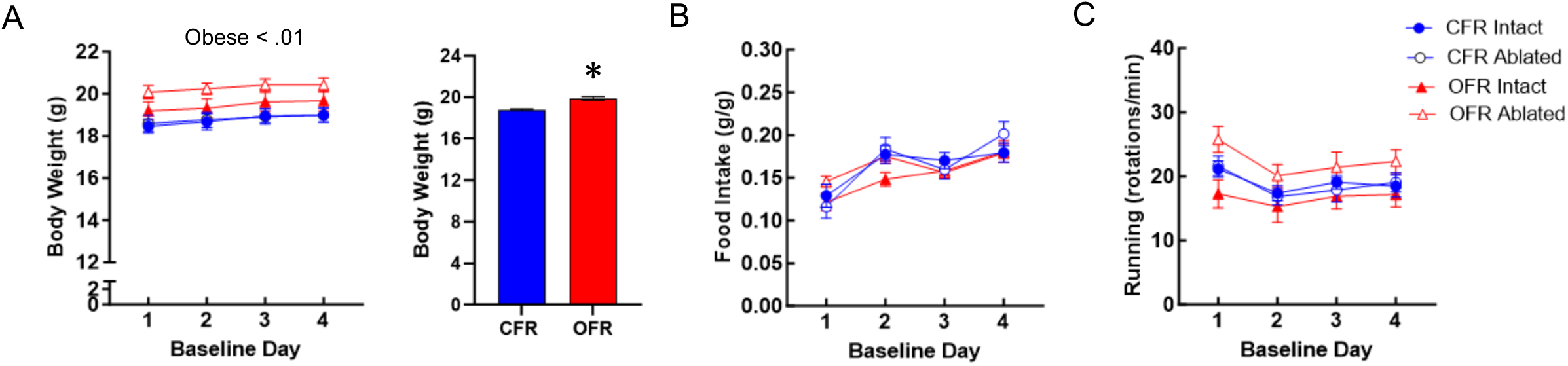
Experiment 2: Effects of obese WAT transplant and neonatal AgRP ablation during baseline. (A) OFR mice weighed more than CFR mice over the 4 days. (B) Food intake in grams divided by body weight of intact or ablated CFR and OFR mice over the 4 days. (C) Wheel running revolutions of intact or ablated CFR and OFR mice over the 4 days. Data are adjusted mean values ± SEM, n = 12-14/group. An asterisk (*) indicates a significant effect of transplant group.

During restriction in Experiment 2, OFR ablated mice showed reduced survival compared to both OFR intact mice (Figure 5a), and CFR ablated mice (Figure 5b), as revealed by a transplant and ablation interaction (Χ^2^=5.40, p<0.05)(Figure 5a,b). Post-hoc hazard ratios indicated that OFR ablated mice dropped out of the paradigm 0.35 times faster than CFR ablated mice (p<0.01)(Figure 5b), and 0.49 times faster than OFR intact mice, as indicated by a strong trend (p=0.05)(Figure 5b)(95% CI: 1.338-5.596). As in Experiment 1, OFR intact mice showed longer survival times than CFR intact mice; however, this effect did not achieve significance in Experiment 2 (Figure 5b), perhaps due to smaller animal numbers. Lastly, neonatal AgRP ablation did not alter survival in CFR mice (Figure 5a).

**Figure 5.**
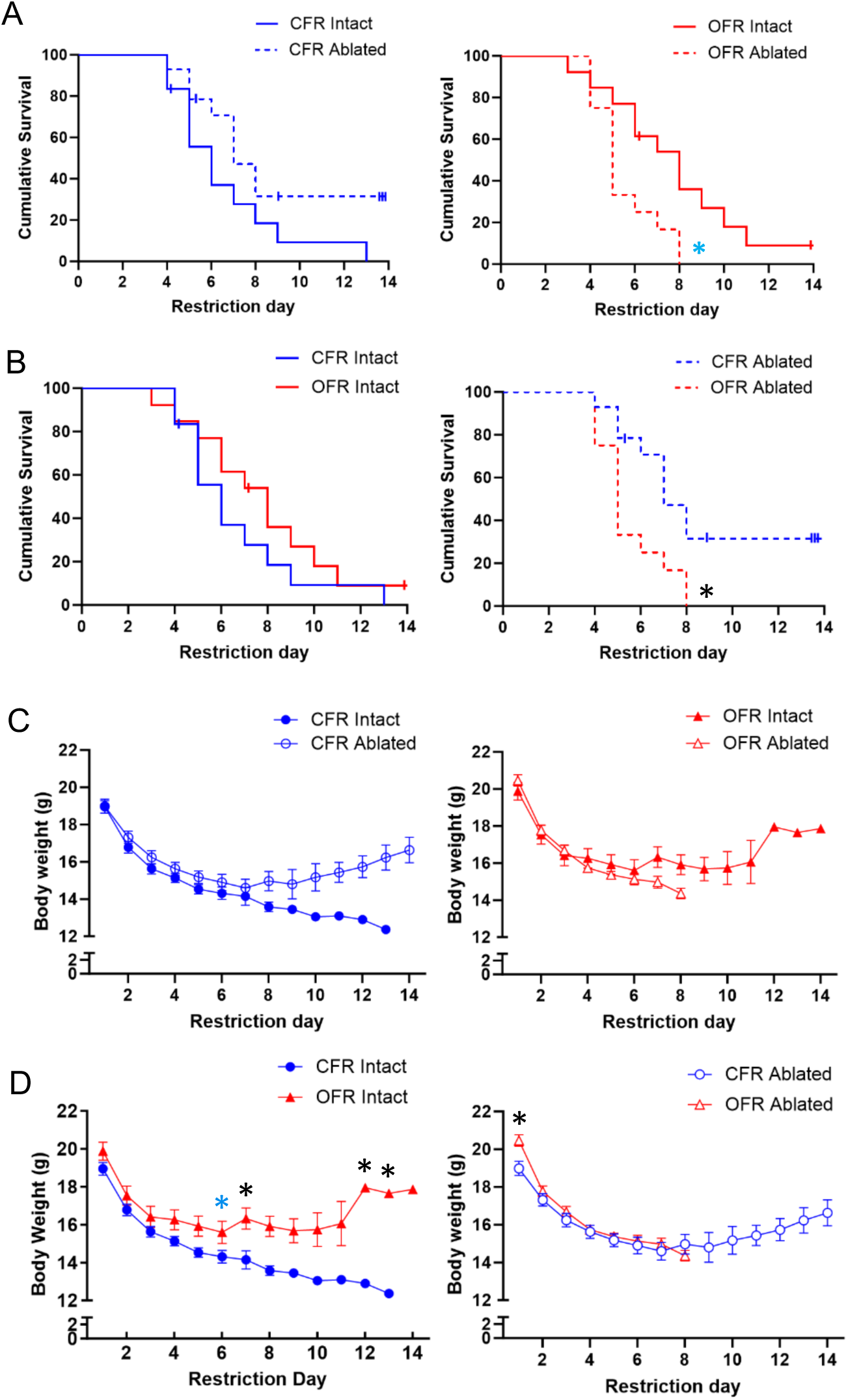
Experiment 2: Effects of obese WAT transplant and neonatal AgRP ablation on survival and body weight during restriction. (A) Ablation reduced survival in OFR, but not CFR mice. (B) Following ablation, OFR mice showed reduced survival compared to CFR mice. (C) No effects of transplant or ablation on body weight. (D) OFR mice showed higher body weight than CFR mice; ablation prevented this effect. Data are adjusted mean values ± SEM, n = 12-14/group. A black asterisk (*) indicates a significant effect of transplant group. A blue asterisk (*) indicates a trend.

A three-way interaction of transplant, ablation, and day was found for body weight during restriction (F(_7,253_)= 2.83, p<0.01; Figure 5c,d). Post-hocs revealed that within intact mice, OFR mice had higher body weights than CFR mice, and this effect reached significance on days 7, 12, and 13 (Figure 5d). Additionally, post-hoc tests indicated a trend on day 6 (p=0.1). To the contrary, within ablated mice, there was no effect of transplant (Figure 5d). Post-hocs also revealed a small increase in body weight in OFR ablated mice relative to CFR ablated mice on day 1 (p<0.05)(Figure 5d). This difference might have resulted from the nonsignificantly higher body weights of OFR ablated mice during baseline (Figure 4a). Although AgRP ablation reduced survival in OFR mice (Figure 5a), ablation did not reduce body weight within CFR mice (Figure 5c). Importantly, in ABA experiments, animals drop out of the paradigm when the weight loss criterion is reached; therefore, some graphed mean values reflect only one animal, such as CFR intact animals (Figure 5c,d) on days 9-12. In sum, obese WAT transplant enhances body weight maintenance in intact recipient mice during ABA, but neonatal AgRP ablation abolishes this effect.

### Effects of WAT transplant and neonatal AgRP ablation on food intake and wheel running during ABA

An interaction of transplant and ablation was found for food intake during restriction (F(_1, 47_)= 5.43, p<0.05; Figure 6a,b). Although post-hoc tests yielded effects and trends for OFR ablated mice to consume less than CFR ablated mice (p=0.06) and OFR intact mice (p<0.05), these raw p values did not withstand correction for multiple corrections. For wheel running, an interaction of transplant, ablation, and day was found during restriction (F(_6, 173_)= 4.09, p<0.001; Figure 6c,d). Post-hoc tests showed that OFR ablated mice ran more than OFR intact mice only on day 2.

**Figure 6.**
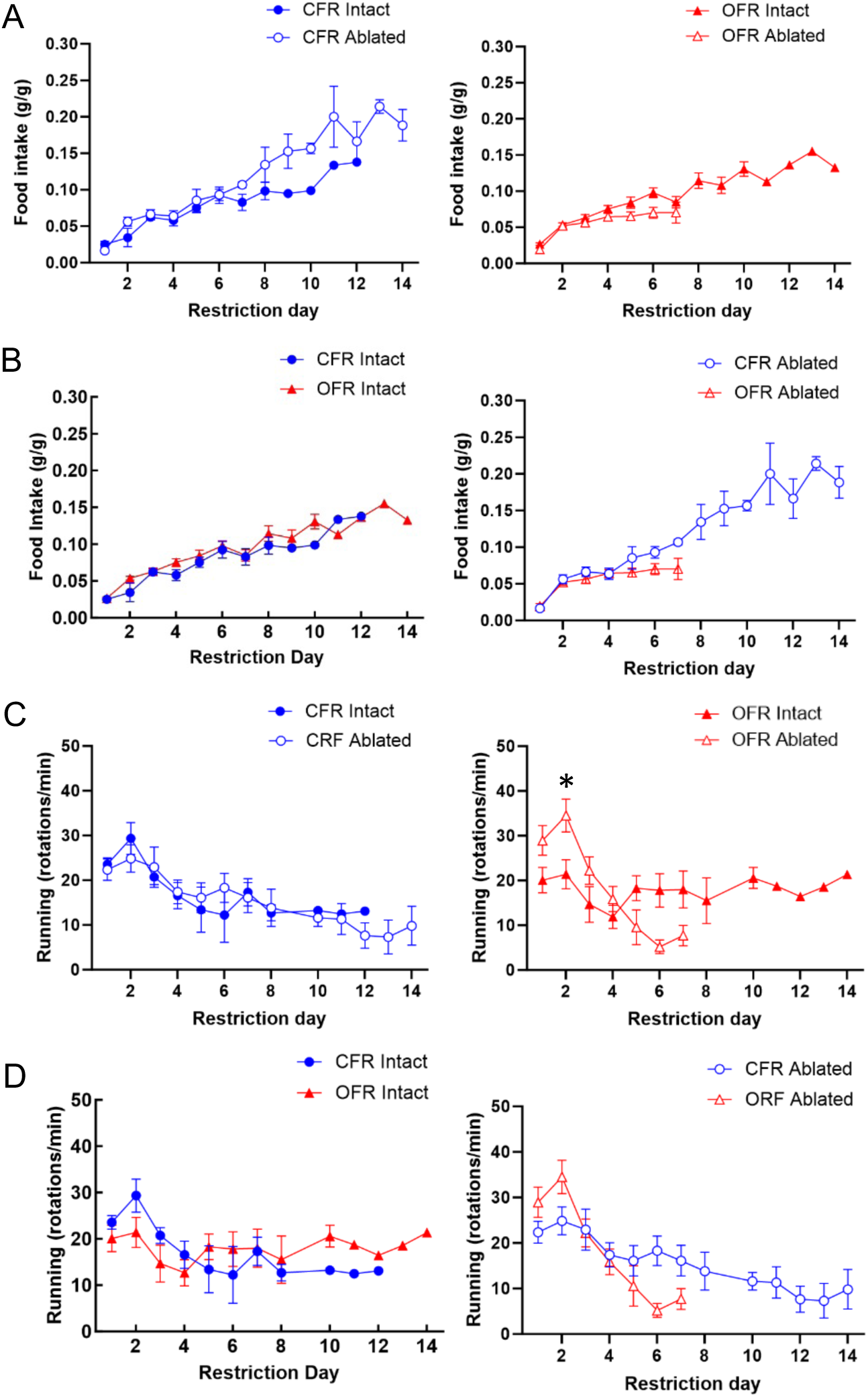
Experiment 2: Effects of obese WAT transplant and neonatal AgRP ablation on food intake and wheel running during restriction. (A) A trend for ablation to reduce food intake was found in OFR, but not CFR mice. (B) Following ablation, OFR ablated mice showed a trend for reduced food intake compared to CFR ablated mice. (C) No overall effect of ablation on wheel running of either transplant group. (D) No overall effects of transplant group within control or ablated groups. Data are adjusted mean values ± SEM, n = 12-14/group. A black asterisk (*) indicates a significant effect of transplant group.

To determine whether transplant or ablation altered wheel running during specific phases of ABA, wheel running during the FAA and PPA periods was also analyzed. No main effects or interactions were found for wheel running during the FAA (Suppl. Figure 2a,b) or PPA (Suppl. Figure 2c,d) periods.

## DISCUSSION

Here we show, for the first time, that transplanting WAT from obese mice into normal recipient mice reduces weight loss of the recipients during the ABA paradigm. We also show that this obese WAT transplant-mediated weight loss resistance during ABA can be abolished by neonatal ablation of AgRP neurons in the recipients. As the largest endocrine tissue of the body, WAT produces and secretes hormones and exosomes containing other active substances into the bloodstream, thus communicating with other tissues in the body, including the hypothalamus (53). Furthermore, recent work suggests that AgRP neurons sense energy loss and drive feeding, running, and fuel mobilization during the ABA paradigm (44). Our present findings indicate that AgRP neurons are required for obese WAT transplant to improve body weight maintenance during ABA, an effect that likely involves communication from obese WAT to AgRP neurons through diffusible factors. Our findings are consistent with those from the largest GWAS of AN to date, which reported significant genetic correlations between AN and metabolic, lipid, and anthropometric traits, suggesting that AN is a metabo-psychiatric disorder (11). Our present findings suggest that diffusible factors released by obese WAT in the periphery could be harnessed as novel therapeutics for AN.

During the 4 day baseline period of the ABA paradigm, we did not observe any consistent effects of WAT transplant or neonatal AgRP ablation. The only exception was a small but significant increase in body weight across baseline days in OFR versus CFR mice. This small effect was observed in Experiment 2 (Figure 3a), but not Experiment 1 (Figure 2a). Thus, obese WAT transplant does not reliably produce increases in body weight under baseline conditions; no other study has examined this effect, to our knowledge. This small difference in baseline body weight in Experiment 2 did not confound our findings regarding body weight during restriction, since OFR ablated mice had the highest baseline body weight, but showed the most rapid dropout from the ABA paradigm (Figure 5a,b). Lastly, our findings are consistent with other reports that neonatal AgRP ablation does not alter body weight or food intake under baseline conditions (44, 51).

During the restriction phase of ABA, OFR mice showed increased survival compared to CFR mice in Experiment 1 (Figure 3d), and increased body weight compared to CFR mice in Experiment 2 (Figure 5d). While the survival measure indicates the day by which mice have lost 25% of their baseline day 4 body weight, the body weight measure indicates the average weight of mice still remaining in the paradigm. Thus, these two measures capture different aspects of body weight loss during the restriction period. The ability of obese WAT transplant to prevent weight loss during restriction was evident in the survival measure in Experiment 1, and in the body weight measure in Experiment 2. In sum, the results of the two studies both indicate that obese WAT transplant reliably reduces weight loss in intact mice during the restriction period of ABA.

One report suggests that normal mice receiving AgRP ablation during the neonatal period show substantial reductions in survival, body weight, food intake, and wheel running during the restriction phase of ABA (44). However, we did not find any effects of neonatal AgRP ablation in CFR mice on any measure during restriction (Figures 5a,c and 6a,c). However, our present studies tested 14 week-old mice using a 3h period of food access during restriction, while previous work tested 5-7 week-old mice using a 2h period of food access (44). These differences in experimental conditions likely account for the discrepancy in results, since longer periods of food availability and the use of older animals both reduce the amount of weight loss during ABA (22, 25). On the other hand, we found that neonatal AgRP ablation reduced body weight measures during restriction in OFR mice, perhaps due to their higher body weights compared to CFR mice. Specifically, ablation reduced survival in OFR mice, compared to both OFR intact mice (a trend, p=0.05) and CFR ablated mice (Figure 4a,b). Similarly, OFR intact mice showed higher body weight than CFR intact mice during restriction (Figure 4d), but neonatal AgRP ablation completely eliminated this effect (Figure 4d). Lastly, a small increase in body weight observed in OFR ablated mice on the first day of restriction likely resulted from the small increase in body weight observed in this group during baseline (Figure 4a). In sum, AgRP neurons are essential for the ability of obese WAT transplant to prevent weight loss during ABA.

Overall, obese WAT transplantation did not alter food intake in the present studies. In Experiment 1, a sharp but transient increase in food intake was observed in CFR mice relative to OFR mice on days 4 and 5 of restriction (Figure 3f); however, food intake rapidly became comparable between the transplant groups on the remaining days. This transient increase in food intake in CFR mice likely emerged due to the rapid drop out of over 60% of CFR mice on days 3 and 4, with few remaining CFR mice having large food intake values. Furthermore, no difference in food intake was found between CFR intact and OFR intact groups in Experiment 2 (Figure 6b). Yet, OFR ablated mice showed a trend to eat less than both OFR intact and CFR ablated mice (Figure 6a,b). Thus, neonatal AgRP ablation might have reduced survival in OFR mice by reducing food intake, at least in part. The observed trend for OFR ablated mice to eat less during restriction is consistent with our previous work showing that obese WAT transplants contain specific factors that can encourage feeding (34).

Neither obese WAT transplantation nor neonatal AgRP ablation altered wheel running during restriction overall, or during the FAA or PPA periods (Supplemental Figures 1 and 2). In Experiment 1, CFR mice showed increased running only on day 6, when few animals remained in the paradigm (Figure 3g). In Experiment 2, only a small increase in running was observed in OFR ablated versus OFR intact mice on day 2 of restriction, which might have resulted from nonsignificant increases in running by this group during baseline (Figure 4c).

Adipose tissue communicates with the brain through factors secreted into the bloodstream (35, 36). Since OFR mice showed increased body weight and survival during ABA without changes in food intake or wheel running, this effect likely resulted from altered metabolism, mediated by obese WAT-secreted factors interacting with AgRP neurons (44, 54, 55). For example, activated AgRP neurons shift metabolism towards energy conservation and lipid storage through several mechanisms including reduction of energy expenditure, elevated carbohydrate utilization, and reduced sympathetic outflow, independent of changes in food intake (37, 46–49). We recently found that control and obese WAT transplants show different patterns of gene expression, which may result in differences in secreted factors (34). Identifying the factors secreted by obese WAT to increase body weight maintenance during ABA may inform novel treatment approaches for AN and related disorders. Lastly, our findings also suggest that autologous WAT transfer following reprogramming to reflect an obese state should be explored as a potential treatment for AN.

The present studies have several limitations. We used female mice as both donors and recipients, since girls and women show far greater vulnerability to developing AN. However, investigations should be extended to males, as boys with AN present with cognitive and behavioral symptoms that are as severe as their female counterparts (56). Furthermore, numerous sex differences have been reported in WAT function in humans, including adipogenesis, the storage and release of fatty acids, and secretory function (57–60). Additionally, neonatal ablation of AgRP neurons induces compensatory changes in networks which may complicate interpretation of results. However, adult AgRP neuron ablation results in death, precluding the use of this manipulation to avoid compensation (51).

In summary, transplanting WAT from an obese donor into a normal weight recipient attenuates weight loss during ABA in the recipient, and neonatal AgRP ablation abolishes this effect. The increased body weight maintenance exhibited by OFR mice is not associated with alterations in feeding or wheel running, but may involve metabolic changes. Findings strongly suggest that an obese WAT transplant communicates with AgRP neurons to promote body weight maintenance during ABA. These findings provide essential groundwork for the identification of obese WAT-derived factors for developing treatments for AN, the deadliest psychiatric disorder.

## ACKNOWLEDGEMENTS

SD, OO, and AL were supported by a Department of Defense Discovery Award (W81XWH2210084) and R21-MH128574, both to SD. RZ was supported by the Larry L. Hillblom Foundation Postdoctoral Fellowship 2019-D-007-FEL. We would like to thank Eeshi Uppalapanti and Imran Kyeseo for assistance with running experiments.

---

**Supplementary Figure 1.**
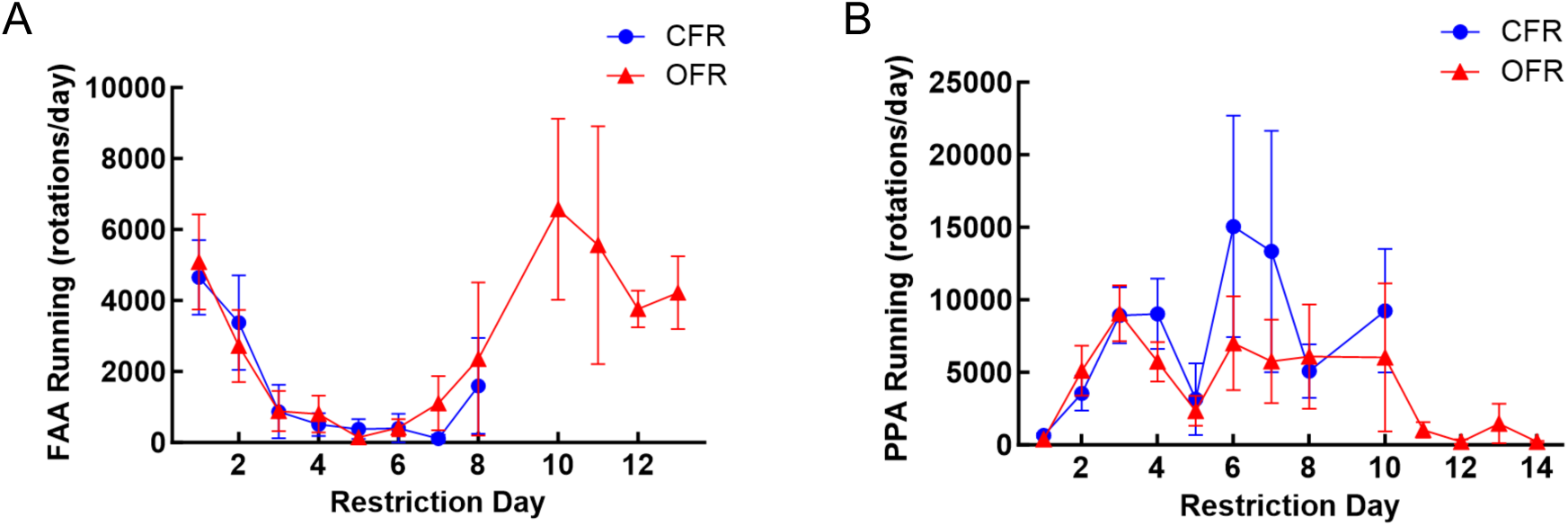
Experiment 1: Transplant had no effect on wheel running during the FAA (A) or PPA (B) period of restriction. Data are adjusted mean values ± SEM, n = 21/group.

**Supplementary Figure 2.**
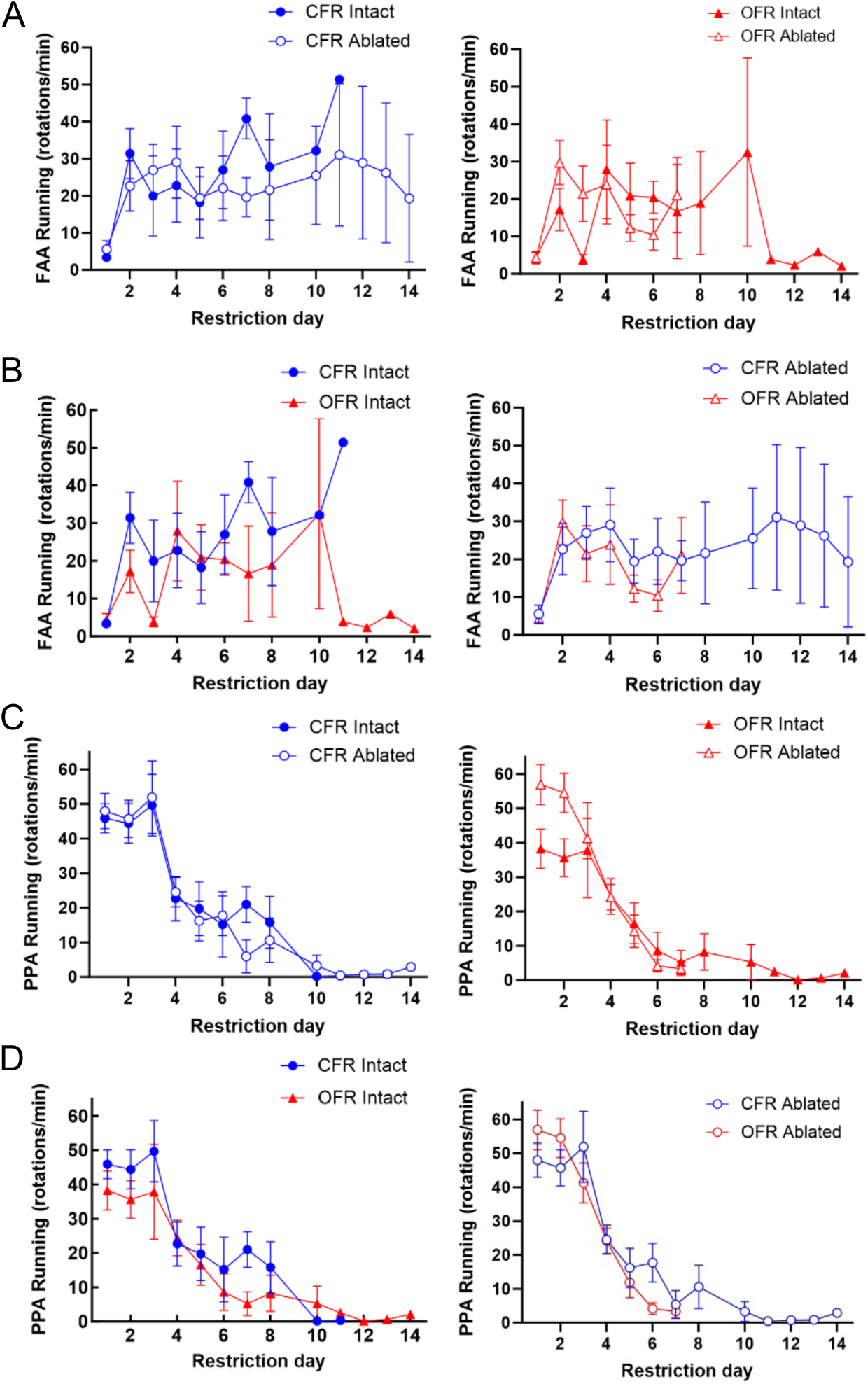
Experiment 2: Neither transplant nor ablation altered wheel running during the FAA (A,B) or PPA (C,D) period of restriction. Data are adjusted mean values ± SEM, n = 12-14/group.

